# Competing females strategically combine signals and physical aggression differently from males in a polygynous tropical lizard

**DOI:** 10.1101/2022.09.27.508996

**Authors:** Devica Ranade, Ratna Karatgi, Shrinidhi Mahishi, Kavita Isvaran

**Affiliations:** Centre for Ecological Sciences, Indian Institute of Science, Bangalore-560012, India; Indian Institute of Science Education and Research (IISER), Pune-411008, India

**Keywords:** Intrasexual competition, females, polygynous mating system, lizards, social selection, sexual selection, Psammophilus dorsalis

## Abstract

Intrasexual competition, the intense competition between individuals of the same sex, is a strong evolutionary force that is well known to select for elaborate and spectacular traits in males. In contrast, female-female competition is poorly understood. Although historically expected to be weak, evidence for female-female competition is fast accumulating. Yet, systematic studies of such competition are rare. Here, we frame and test for a strategy of competition in females in a wild population of a classically polygynous lizard, *Psammophilus dorsalis*. We used models of female *P. dorsalis* to simulate a gradient in perceived competitor threat to individually tagged, wild females on their territories, and comprehensively measured their signalling and aggression responses. Similar experiments were performed with males. Our results clearly demonstrate a complex, threat-sensitive strategy of direct competition among females in this species with conventional sex roles. Females systematically escalate their use of diverse signals along a gradient in threat and show surprisingly high levels of physical aggression. Comparisons with the responses of males to simulated threats reveal distinct sex differences in competition strategies that match theoretical expectations derived from life history. Using these findings, we argue that female-female competition may be complex but cryptic, requiring experimental manipulations to uncover.

## Introduction

Intense competition between individuals of the same sex strongly selects for striking and elaborate traits. Such intrasexual competition has largely been studied in males because males more frequently possess exaggerated traits than do females. Such exaggeration is thought to result from the competition for mates, which is theoretically expected to be stronger among males than among females (Darwin 1871; Shuster & Wade 2003). However, interest in intrasexual competition in females is now rapidly growing, driven by a revived recognition that females do compete among themselves, but more commonly for breeding opportunities and resources that enhance offspring reproductive value than for mates (West-Eberhard 1979, 2014; Rosvall 2011; Stockley & Bro-Jørgensen 2011; Tobias *et al*. 2012; Clutton-Brock & Huchard 2013b, a; Stockley & Campbell 2013; Hare & Simmons 2019). Evidence for female-female competition is accumulating from diverse taxa (Rosvall 2011; Tobias *et al*. 2012; Stockley & Campbell 2013). But what general form should female-female competition take and what kinds of traits should therefore evolve? Strategies of female-female competition are well known from rare societies, such as sex-role reversed species (Clutton-Brock 2007, 2009; Geberzahn *et al*. 2009), eusocial insects (Ratnieks *et al*. 2006; Bhadra *et al*. 2010) and cooperatively breeding species (Young *et al*. 2006; Clutton-Brock 2007; Cant & Young 2013). However, we lack an understanding of how females compete in more common societies like socially polygynous systems in which female competition is typically believed to be weak. Here, we build a framework for understanding strategies of competition in females and test our predictions in a socially polygynous system.

Regardless of what females are competing for, they rarely experience immediate fitness benefits from signalling and contests because they must still survive to produce offspring and, in some cases, provide maternal care. Additionally, they may face large costs: conspicuous or energetically expensive behaviours and aggression may increase their risk of dying before they produce offspring or may affect their investment in offspring (Stockley & Bro-Jørgensen 2011; Clutton-Brock & Huchard 2013a). In contrast, males competing for fertilizations can gain fitness relatively quickly, especially in polygynous species with limited paternal care. Furthermore, injured males are known to mate, and given their lower investment in offspring, they do not experience the long-term costs that females do. These differences in benefits and costs of competition lead us to expect that costly signalling and physical aggression should not be commonly seen in females. Rather, females should signal using relatively inconspicuous traits, be keenly sensitive to the level of competitor threat, and escalate to costly signalling and aggression only when the threat is high.

We tested this hypothesis that females should display complex, threat-sensitive competition strategies using Peninsular rock agamas, *Psammophilus dorsalis*, in their natural habitat. In this socially polygynous species, males are larger and more conspicuously coloured than females. Adults typically survive through only one breeding season (Deodhar & Isvaran 2017). Males and females use diverse signals, which they broadcast to the larger neighbourhood, or direct toward individuals in close encounters. We predicted that during typical broadcast signalling (in the absence of immediate competition), females should signal relatively inconspicuously. We manipulated perceived competitor threat through simulated intrusions and predicted that females should intensify their signalling and escalate to overt aggression only under high levels of threat. We show that females display a complex, threat-sensitive strategy of competition, which incorporates multiple signals and aggression. By carrying out similar field experiments on males, we demonstrate distinct sex differences in competition strategies. We use these findings to argue that female competition may be widespread and complex, but cryptic, requiring observations of subtle signalling behaviours and manipulations to uncover.

## Materials and Methods

### A. Study species and study site

*Psammophilus dorsalis* is an agamid lizard found in peninsular India (Smith 1935; Daniel 2002). The study was conducted in the hills of the Eastern Ghats in Andhra Pradesh, India (13°32’ N, 78°28’ E). The habitat consists of open spans of bare granite rocks, or sheet rocks, surrounded by scrub and deciduous vegetation growth. Individuals are largely annual, and the mating season lasts from May-October; offspring appear from August-February and mature into adults the following year (Deodhar & Isvaran 2017) Males are larger and more brightly coloured (Deodhar & Isvaran 2017). Such dimorphism is typical of a classical socially polygynous mating system, where intrasexual competition is expected to be strong in males but weak in females. Both sexes defend territories from same-sex individuals in the mating season. A breeding male’s territory typically overlaps multiple female territories (Ranade & Isvaran 2022).

Both sexes display using a set of signals, which includes stereotyped movements like head bobs and push ups, and postures like tail raise, gular extension, and gape (Figure 1 Additionally, they show context-dependent dynamic colour change (Batabyal & Thaker 2017; Deodhar & Isvaran 2018). In the mating season, males are usually observed in yellow-black colouration or less frequently in pale yellow-black (Deodhar & Isvaran 2017, 2018) (Figure 1). However, the colour can change within seconds to orange-black while courting a female; or to red-yellow during competitive encounters (Figure 1) (Deodhar & Isvaran 2018). Breeding females are usually drab grey in colour with an off-white head (Figure 1) but the colour can change dynamically to grey body-orange head or to dark body-orange head (Figure 1).

**Figure 1:**
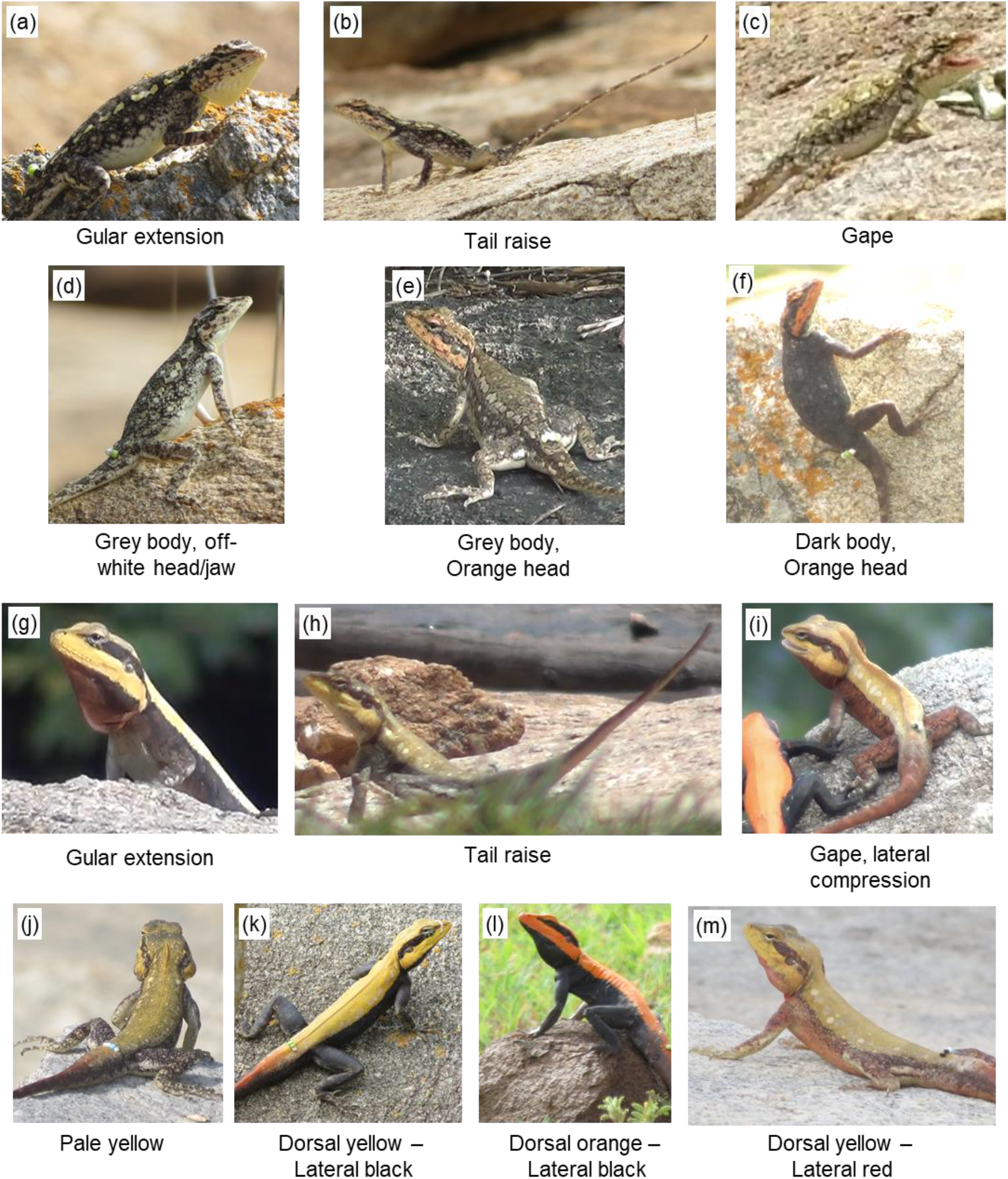
Pictures representing postures (a,b,c, g,h,i) and colours (d,e,f,j,k,l,m) exhibited by females (a-f) and males (g-m)

### B. Data collection

To test for strategies in intrasexual competition in females, we examined female signalling and aggression along a gradient in competitor threat: (i) the absence of immediate threat (unmanipulated, broadcast signalling), and (ii) low and (iii) high levels of (simulated) competitor threat. We also studied male behaviour along a similar gradient, to compare female strategies with those of the well-studied male strategies in socially polygynous societies.

*P. dorsalis* individuals spend most of their time perched on rocks or boulders (henceforth perches). The territory of every individual has at least one of these perches (Radder *et al*. 2006; Ranade & Isvaran 2022). Territory holders are expected to experience a greater threat from intruders who are on their perch than those who are off the perch on a peripheral part of the territory. To simulate high and low intensity immediate competitor threats, life like 3D printed models were presented to individuals ‘on’ their regularly used perch (henceforth on-perch) and 1 metre away from their regularly used perch but within their territory (henceforth off-perch).

Data was collected in the mating seasons (May-October) of 2016 and 2017 between 0700 and 1100 hours in the morning and 1600 and 1800 hours in the evening, since this is when lizards are most active. Focal observations (Altmann 1974) of individuals were made by video recording them. First, the behaviour of a randomly chosen focal individual was video recorded for 30 minutes in its natural environmental conditions, when an immediate intruder threat was not experimentally simulated, to measure its broadcast or baseline signalling intensity. After the 30 minute observation, a 3D printed, static, life-like, size-matched model (Supplementary Figure S1) was presented to the focal individual either ‘on’ or ‘off’ its perch (details of models used in Supplementary material). Focal individuals typically went inside a refuge when the observers approached to place the model at the appropriate position, but emerged within 1-15 minutes of model presentation.

Quantification of behaviours was started only after the individual emerged from the refuge and was judged to be in direct line of sight of the lizard model. Quantification was continued for 20 minutes thereafter. Additionally, because the observations were made in their natural environment, the time for which other males and females (henceforth audience) were present within 5m of the focal individual was always recorded.

Model presentation trials were conducted on 24 females and 22 males. Each randomly selected individual received both treatments (on/off perch), 3-10 days apart. The sequence of treatments was randomised (further details of methods in the supplementary material) The videos were transcribed using BORIS (Behavioural Observation Research Interactive Software) version 2.72 (Friard & Gamba 2016). Signals were classified into body movements (head bobs, push ups), body postures (gular extension, tail raise, gape, lateral compression (shown by males only) and time spent in different colours. Rates of body movements and proportion of time spent in different postures and colours (Figure 1) were calculated for every focal observation session. The number of aggression events without contact (chases) and with contact (physical fights) were also noted.

### C. Statistical analysis

Statistical analysis was conducted using the statistical software RStudio (R Core Team 2014) version 1.2.1335. Please note that multiple null hypothesis significance tests are reported, however, multiple p-value adjustments have not been performed, because its use is still debated (e.g., Moran 2003). Instead, we base our inferences on the larger concordance of results than on individual p-values.

#### C.i. Signalling and aggression

First, to test the prediction that during broadcast signalling, females signal with less conspicuous signals, more inconspicuous than those in males, each signal was modelled as a function of sex and proportion of time for which male and female audience were present within 5 m of the focal individual during a focal observation (to control for varying audience). Only signals that were shown by both sexes were analysed. Generalised linear mixed effects models (GLMMs) with negative binomial error structure (to account for overdispersion) and a log link were run for signals represented as counts (head bobs and push ups) using the function ‘glmmTMB’ (Brooks *et al*. 2017). The time for which an individual was in sight during a focal observation was included as an offset term to account for variation in sampling effort. Identity of an individual was included as a random effect. For signals represented as proportion of time spent in a posture (gular extension, tail raise) colour state, linear mixed effects models (LMEs) were run with the same fixed and random effects as in the GLMMs. Gape was excluded from this analysis as it was rarely recorded (less than 5% of observations). Additionally, because males and females show different colour states, colours were divided into relatively camouflaging (in females: grey body-off white head; in males: dorsal pale yellow-lateral light black) and relatively conspicuous (in females: grey body-orange head and dark body-orange head; in males: dorsal yellow-lateral black, dorsal orange-lateral black and dorsal yellow-lateral red).

Next, to test the prediction that females should be sensitive to the level of intruder threat, even more than males, each signal that was shown by both males and females was modelled as a function of the type of threat experienced, sex and their interaction. Threat type consisted of three levels; no (broadcast signalling), low (off-perch model presentation), and high (on-perch model presentation). The interaction term was included to model the possibility that males and females use signals differently when responding to a competitor. As before, for response variables that were counts (head bobs, push ups), GLMMs with a negative binomial error structure and a log link were run. The duration for which the lizard was in sight during a focal observation was included as an offset term and individual identity as a random effect. LMEs with similar fixed and random effects as the GLMMs just described were run on the proportion of time spent in postures (gular extension, tail raise, gape). Postures and colours shown by one sex only were modelled as a function of treatment type using LMEs with individual identity as a random effect. For all models, permutation tests were run and 95% bootstrapped confidence intervals on model coefficients were calculated to increase our confidence in models (further analysis details in supplementary material).

The number of focal observation sessions in which contact aggressive behaviours occurred was compared between treatment types and sexes using 95% confidence intervals.

#### C.ii. Complexity in female competition strategies

The complexity of response to the level of threat was examined in detail. First, as described above, we quantified the frequency of body movements and time spent in posture and colour states at varying levels of competitor threat. Second, we counted the number of different signals displayed by individuals during a focal observation session, for the different treatments. Third, we identified the sequence in which different signals were first displayed by every focal individual during a focal observation session. Please note that we did not examine all shifts from one signal to the next but focussed on the sequence in which signals appeared for the first time in a focal observation session. Additionally, since our interest lay in signals used in direct competition, only signals which appeared to vary across different levels of competitor threat were included while working on points 2 and 3.

## Results

First, we examined the most common signalling context, non-directed or broadcast signalling, where an individual signals from within its territory to the larger neighbourhood and there is no immediate intruder threat. As predicted, females used comparatively inconspicuous signals during broadcast signalling, more so than did males. Females used relatively inconspicuous head-bobs more frequently than the more conspicuous push-ups (Figure 2a, Supplementary Table S1). Males showed a similar trend. However, females used push ups less frequently than did males (Figure 2a, Supplementary Table S1). Furthermore, females spent much more time (*ca.* 96% more) in camouflaging colour than in conspicuous colour. In contrast, males spent *ca.* 73% more time in conspicuous than in camouflaging colour (Figure 2c, Supplementary Table S1). Additionally, compared with males, females spent much less time in posture with extended gular and more time in a posture with raised tail (Figure 2b, Supplementary Table S1). Neither sex showed aggression involving physical contact, i.e., physical fights, suggesting that it is rare. When a same-sex intruder was detected within 2 metres of the focal individual within its territory (females n = 2, males n = 3), the resident chased away the intruder.

**Figure 2:**
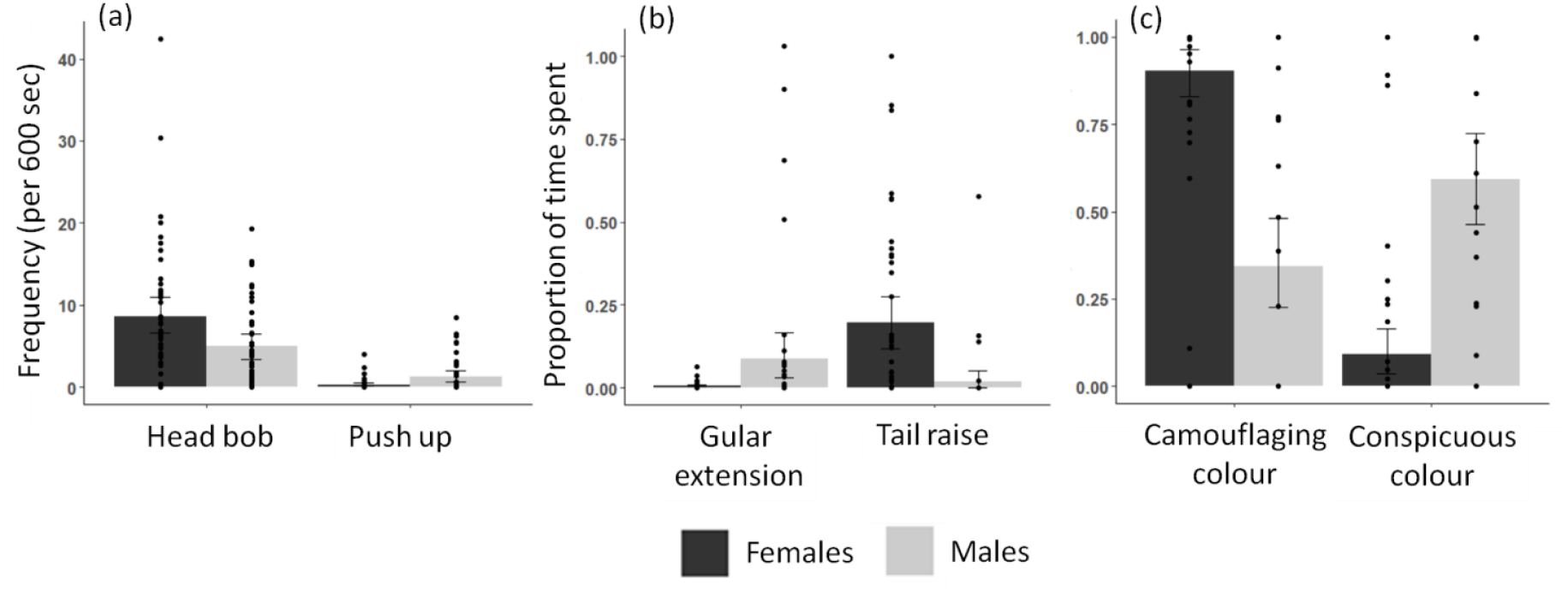
Broadcast signalling in females and its comparison with males. Rates of body movements (a) and time spent in posture (b) and colour states (c) in the absence of an immediate intruder threat by females (black bars) and males (grey bars). Error bars represent 95% bootstrapped confidence intervals around the mean and black dots represent raw data points. Females appear to signal using relatively less conspicuous head bobs more often than the more noticeable push ups, more so than males (a). Females spend more time in a raised tail posture and less time in extended gular posture as compared to males (b). Females were observed in camouflaging colour more often than in conspicuous colour. In contrast, males were observed more often in conspicuous colour (c). Statistical tests supporting results are reported in Supplementary Table S1.

Next, we examined female signalling and aggression at varying levels of competitor threat, by simulating intruder threat using 3D printed models. Females showed a striking response to increasing competitor threat (Figure 3). They substantially increased their use of diverse signals. The frequency of head bobs and push ups and the duration of display behaviours like gular extension and gape showed clear increases with increasing threat (statistical support shown in Supplementary material table S2). Males also responded to threat, but there were clear sex differences (Figure 3, Supplementary material table S2, S3). While males showed increases similar to those of females in some signalling behaviours (e.g., gape, gular extension), they showed no clear increase in other behaviours in contrast to the pattern in females (head bobs and push ups) (Figure 4, Supplementary Table S2). In addition, they increased their use of some signals that were never seen in females including a posture (lateral compression), and a colour signal (dorsal yellow-lateral red) (Figure 4, Supplementary Table S3). In contrast, unlike males, females did not appear to use colour signals in direct competition (Figure 4, Supplementary Table S3).

**Figure 3:**
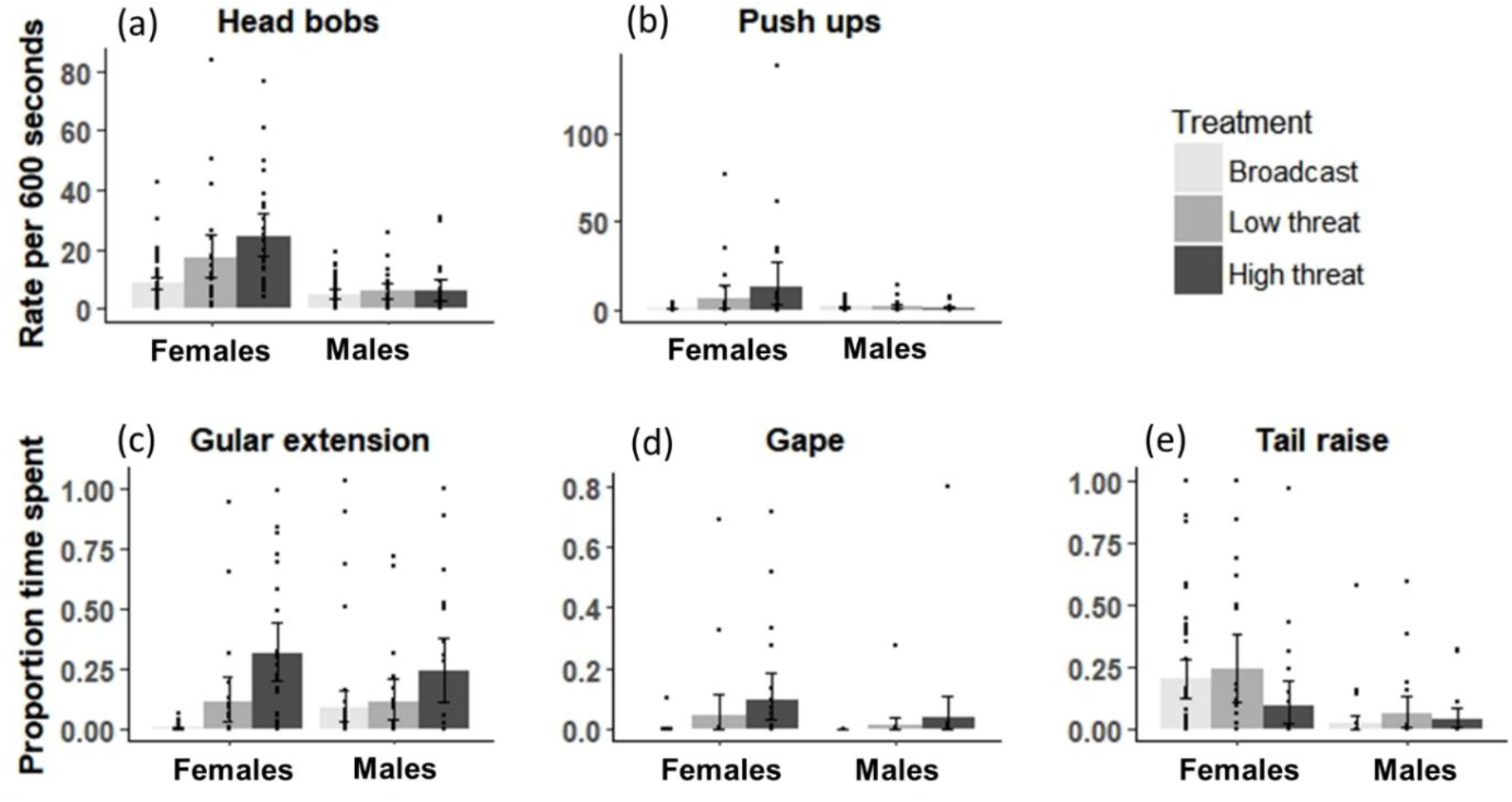
Comparison between frequencies of head bob (a) and push up (b) and proportion of time spent in gular extension (c), gape (d) and tail raise (e) with changing level of intruder threat in females and their comparison with males. Error bars represent 95% bootstrapped confidence intervals around the mean and black dots represent raw data points. Females increased number of head bobs and push ups with increasing level of threat, unlike males. Females, like males, increased time spent in gular extension and gape with level of intruder threat. Tail raise did not seem to be a response to intruder presence in either sex. Statistical tests supporting relationships are reported in Supplementary Table S2.

**Figure 4:**
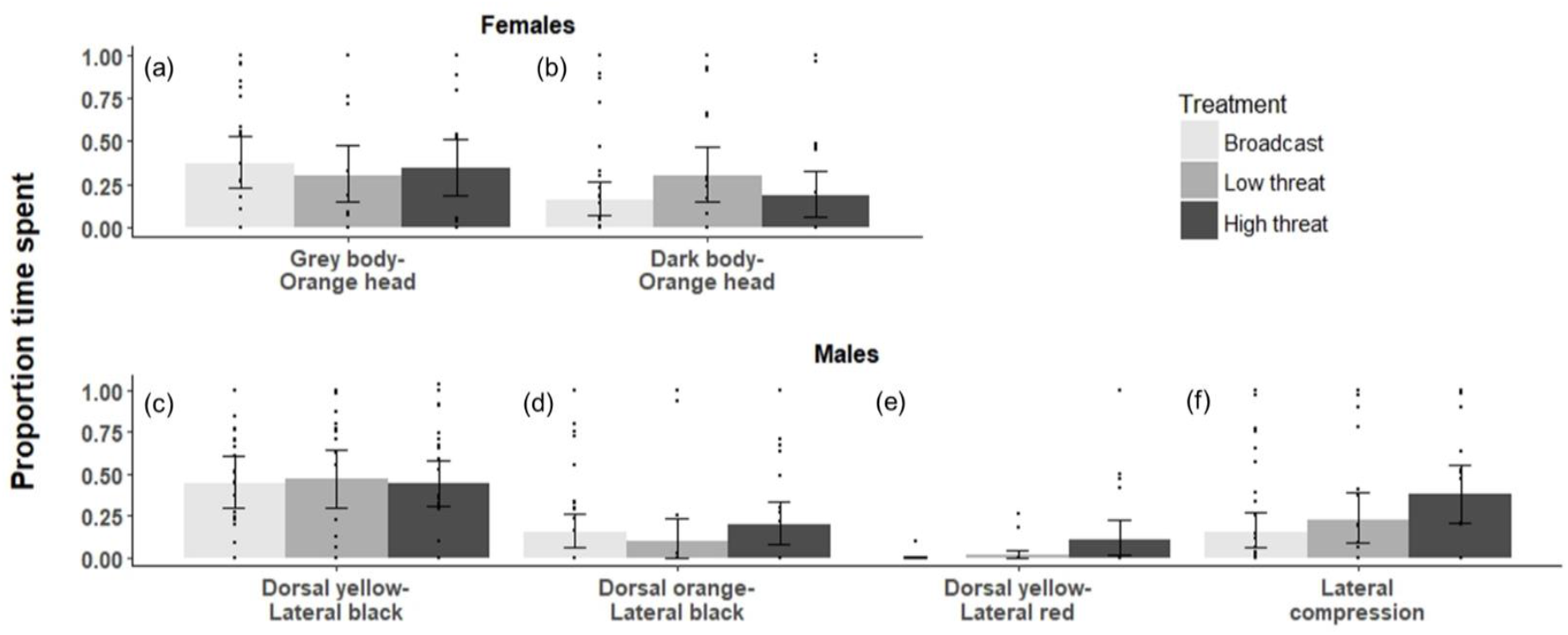
Comparisons in proportion of time spent in colours (a,b,c,d,e) and posture (f) unique to one sex. Error bars represent 95% bootstrapped confidence intervals, while the dots represent the actual data points. Time spent by males in laterally compressed body posture and dorsal yellow-lateral red colour increased with increase in level of intruder threat. Statistical tests supporting results are reported in Supplementary Table S3.

Note that we also confirmed through control model presentations that 3D printed lizards used in the experiments were perceived as competitor lizards and not as novel objects (please see supplementary material for control model experiments).

Next, we analysed if females increase aggression in response to varying levels of intruder threat and if this differs from males. Interestingly, females showed a dramatic increase in aggression, through biting or pushing the simulated intruder, in response to level of threat. First, in the broadcast signalling context, intrusions by other individuals into a resident’s territory were rare; accordingly, aggression was also rare. Next, among 24 females tested with both low and high levels of intruder threat, a striking 11 (46%) females showed aggression towards intruders on an attractive part of the territory (high threat) while only 2 (8%) females were aggressive towards intruders posing a lower level of threat. In contrast, out of 22 males, only 2 (9%) and 1 (4.5%) males were aggressive towards high and low threat competitors respectively (Figure 5).

**Figure 5:**
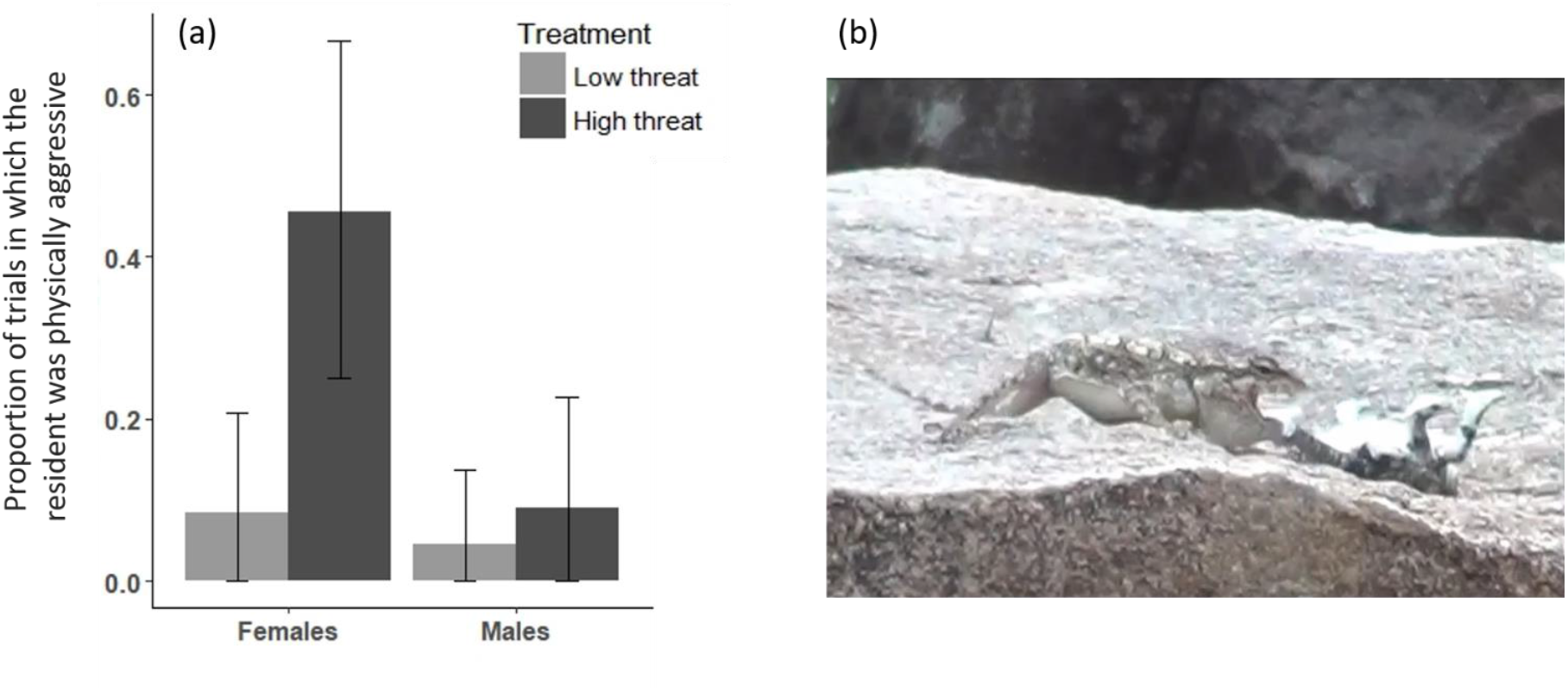
(a) Proportion of times resident females were physically aggressive with the simulated intruder at varying levels of threat and their comparison with males. Error bars represent 95% bootstrapped confidence intervals. Females were strikingly more aggressive at high level of intruder threat. (b) Photograph of a resident female being aggressive with a simulated intruder (3D printed lizard model)

Examining signalling and physical aggression in detail clearly indicated that females show a complex, threat-sensitive strategy of competition, which involves multiple signals and aggression. First, as we describe above, the intensity of individual displays (i.e., higher rates of displays and proportions of time spent) increased along the gradient in competitor threat. Second, the number of unique signals used by females also increased with increasing threat: from 1.5 ± 0.1 (mean ± se) unique signals during broadcast signalling, to 1.9 ± 0.2 signals during simulated low immediate intruder threat, and 2.9 ± 0.2 signals during high immediate intruder threat. Third, these different signals were strikingly used in a systematic sequence in the presence of an immediate intruder threat, providing information on how females escalate in the face of a threat. We examined the order in which signals were displayed for the first time in a focal session. In the absence of an immediate intruder, that is, during broadcast signalling, female signalling was limited to head bobs with only rare instances of other signals. At low and high levels of immediate intruder threat (simulated intruders at less and more attractive areas on the territory), females typically began signalling with head bobs and then used gular extension, following which push ups and gape were used (Figure 6). The sequence was particularly clear in the high threat treatment. Furthermore, in the high threat treatment, either gape or push ups was followed by physical aggression 63% of the time. It is thus evident that females typically do not attack the model directly but use a sequence of signals before attacking the model.

**Figure 6:**
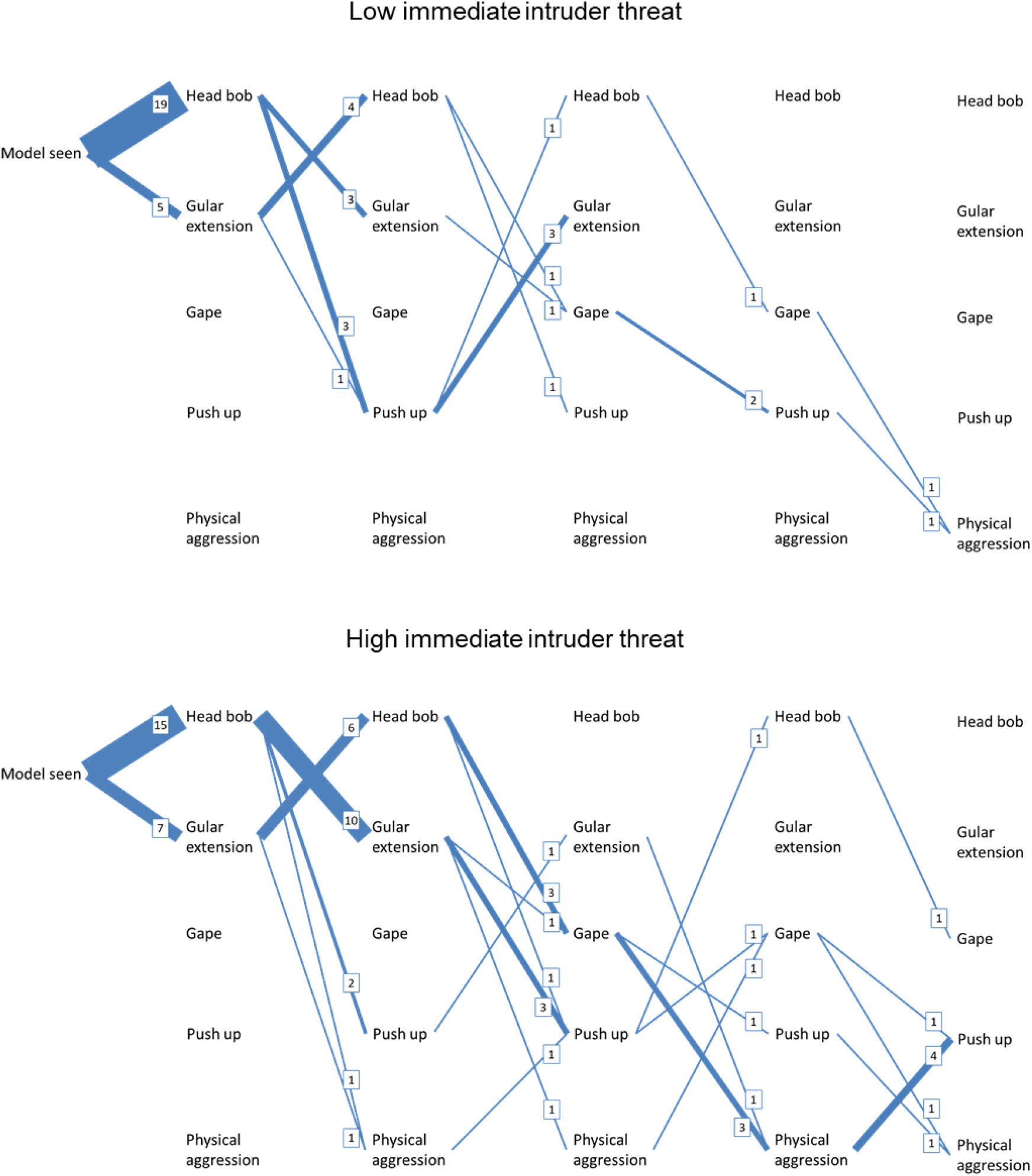
Sequences in which signals and aggression first occur on simulation of low and high immediate competitor threat in females. Each blue line represents only the first shift from one signal to the other during an observation. The thickness of the lines along with the numbers represent the number of times a particular shift has occurred. Females display using a greater number of behaviours at high immediate intruder threat than at low immediate intruder threat (Number of blue lines (shifts) more at high than low intruder threat level). Also, females appear to use a sequence of signals before attacking the model. Please note that only 22 observations were used for understanding sequences at high intruder threat, because we could not video record initial responses of 2 females and were hence uncertain about the sequence of signals first used.

## Discussion

Through observations and field experiments on a wild lizard population, we demonstrate that females in a socially polygynous system show a complex and nuanced strategy of female-female competition that includes both signalling and surprisingly high levels of physical aggression in certain threat contexts. As predicted, when normally signalling to the larger neighbourhood (broadcast signalling in the absence of immediate intruders), females used relatively few and inconspicuous signalling behaviours. Results also supported our second prediction that female signalling should be sensitive to the level of threat. Female signalling increased in complexity along the gradient in competitor threat. As the level of intruder threat increased, females substantially increased the intensity with which they used multiple signals. Females also increased the number of unique signals and displayed them in a characteristic order. Importantly, under the specific context of high immediate intruder threat, females commonly escalated to physical aggression, usually after displaying with a systematic sequence of signals. Comparisons with the responses of males to a similar threat gradient reveal distinct sex differences in competition strategies that match theoretical expectations derived from life history. Taken together, these results on signalling and aggression indicate that intrasexual competition in females is cryptic, but complex and sensitive to the level of competitor threat. To the best of our knowledge, this study is one of the first comprehensive evaluations of strategies of direct competition in females of a species with conventional sex roles, which examines the complete signalling repertoire and aggression, and assesses how females use signals at varying competitor threat levels and when such competition can escalate to aggression.

Female *P. dorsalis* could either be competing for mates or for resources other than mates that are important for survival and production of offspring. A female has access to 2-6 unique males over her lifetime and would have to travel outside her home range to access additional males (Ranade & Isvaran 2022). Therefore, females might compete to gain territories within the home range of high quality males. However, indirect evidence does not support this explanation because females do not change their territories over their lifetime but males do, suggesting that males track female dispersion rather than vice versa (Ranade & Isvaran 2022). Hence, the strong direct competition in females is unlikely to be primarily for mates. Instead, it is likely that females compete for high quality rock perches. These rock perches are expected to provide females with important resources like refuge sites under/between rock crevices for protection against predators, basking sites with good visibility, and foraging sites. Additional work is needed for understanding, in detail, the resources that females compete over. In many other systems too it appears that females compete for resources related to offspring production and survival rather than for mates (West-Eberhard 1979; Clutton-Brock 2009).

The spatial dispersion of *P. dorsalis* suggests that competition among females is resolved in multiple ways. Like many other lizards (Haenel *et al*. 2003; Robles & Halloy 2009) *P. dorsalis* females are solitary and territorial, which is a likely way of avoiding competitors and recurrent contests. During our focal observations, we did not see two or more females on the same perch or in a territory, suggesting that competition is typically resolved through covert means. However, when a female intruder particularly advances towards their perch, resident females signal to and chase away such intruders. Confirming these observations, when the perch of a female, which is potentially the resource of the highest importance was under threat through simulated same-sex intruders, females strikingly increased competition and physically engaged in a fight by biting, pushing and pouncing on the model. These findings suggest that the seemingly docile females can aggressively compete when the threat to resources increases.

The results of our study clearly indicate sex differences in strategies of direct competition. Females used relatively inconspicuous signals under the normal broadcast signalling context, unlike males who used relatively conspicuous signals, especially conspicuous colour patterns. In response to simulated threat too, males but not females used dynamic colour signals. Females, however, showed a more sensitive response to the gradient in threat than did males. They strongly increased their use of multiple behavioural signals (head bobs, push ups, gular extension, gape) with increasing threat level, more so than did males. Furthermore, while both sexes showed little aggression at low threat levels, females but not males dramatically increased their aggression at high threat levels.

These sex differences in competitive behaviour align well with expected sex differences in costs and benefits of competition in a socially polygynous mating system. In this mating system, females are unlikely to experience immediate benefits and are likely to face large costs to competitive traits (Stockley & Bro-Jørgensen 2011; Clutton-Brock & Huchard 2013a). Males, in contrast, can gain large immediate benefits from competition, in the form of mates, especially in species like *P. dorsalis,* where males typically live to experience only one mating system and do not show parental care. Female *P. dorsalis* experience a longer breeding season than do males and can lay multiple clutches (egg laying continues till February while the mating season ends by October). Therefore, while males may benefit from signalling conspicuously even in normal broadcast signalling contexts, females should tailor their costly signalling to the level of threat. They should display costly competitive traits only when the threat is high because their benefits from competition are gained over the long-term while costs can impact egg-laying. An alternative explanation for the sex differences in competitive behaviour is that given the differences in the biology of males and females (for example, larger home range area of males and presence of multiple perches in male home ranges), it is possible that the on-perch and off-perch treatments did not represent the same levels of simulated intruder threat for males and females. Our motivation to carry out the same set of experiments on both sexes was to compare the competition strategy of females with that of males in whom such strategies are well studied. By performing the same set of experiments, we have tried to standardise the presentation of levels of competitor threat. Our study provides a first-level view of how strategies of direct competition may differ between the sexes and suggests future directions for the study of sex differences in intrasexual competition.

Here, we describe a distinct strategy of direct competition in females in a polygynous mating system, in which females escalate their use of multiple signals along a gradient in threat and commonly show physical aggression but only at high levels of threat. Such competition strategies in females have been previously reported only in certain special systems, like cooperative breeding societies and eusocial insects, in which females compete for breeding positions (e.g., Clutton-Brock *et al*. 2006; Bhadra *et al*. 2010). Among common mating systems, such as socially polygynous, socially monogamous, and multi-male-multi-female systems, female competition is now widely reported but systematic studies of strategies of competition are few. Female-female competition has been reported in diverse taxa, including insects (e.g., Shelly 1999; Eggert *et al*. 2008; Papadopoulos *et al*. 2009), crustaceans (e.g., Hughes 1996), arachnids (e.g., Elias *et al*. 2010), fish (e.g., (Kokita 2002; Draud *et al*. 2004), amphibians (e.g., (Summers 1989), reptiles (e.g., Baird & Sloan 2003; Schofield *et al*. 2007; While *et al*. 2009), birds (e.g., Diniz *et al*. 2019; Krieg & Getty 2020) and mammals (e.g., Scott *et al*. 2005; Archie *et al*. 2006). Competition has been described in diverse social contexts, including populations in which females are solitary and territorial (e.g., Hughes 1996), and those in which females live in groups (e.g., Baniel *et al*. 2018). Many studies describe physical aggression among females (e.g., Johnson 1988; Shelly 1999; Bebié & McElligott 2006). Most of these studies report the circumstances under which aggression in females was observed (e.g., depending on breeding status, availability of food, proximity of competitor to nest). But only a few studies, through observations and experiments in the wild and in laboratory conditions, elaborately describe the escalation of competition from signalling to aggression (e.g., Schofield *et al*. 2007; Elias *et al*. 2010) or how females compete differently from males (Hughes 1996; Reece *et al*. 2007; Stuart-smith *et al*. 2007; Elias *et al*. 2010). Studies of how signals and contact aggression are deployed when the degree of threat to the resource varies are scarce.

Thus, systematic studies that comprehensively cover traits used in female-female competition, such as signalling and aggression, along a gradient in threat are still rare. We suggest that this might be because competition between females is likely to be cryptic (as predicted by the nature of costs and benefits of such competition for females). For example, if females typically engage in fairly inconspicuous signalling and escalate to high levels of signalling and aggression only under high levels of threat (as we report in our study), such behaviour may go unnoticed. Females may further try to avoid encountering high levels of threat by, for instance, living in exclusive territories and defending territories using cryptic signals which advertise their ability to defend their territory, resulting in even fewer observations of conspicuous aggression. Thus, intrasexual competition may substantially shape female ecology and behaviour, but remain cryptic. Consequently, such systems may not be chosen for study by researchers interested in intrasexual competition. Therefore, to gain a true picture of female-female competition, we require experimental manipulations on wild populations and ideally along gradients in threat.

In conclusion, we use field experiments in a wild lizard population simulating a gradient in competitor threat to provide evidence for a complex strategy of direct competition in females. Females use diverse visual signals that they tailor in response to the level of threat, and engage in costly physical aggression only after performing a series of signals. We also provide evidence suggesting that females can be even more competitive than males, but only when the resources they are guarding are challenged adequately. These findings from a socially polygynous system highlight that female competition may be widespread and complex, but cryptic, requiring observations of subtle signalling behaviours and manipulations to uncover.

## Acknowledgements

We wish to thank IISc for providing institutional facilities and Rishi Valley school for logistic support during data collection. We are grateful towards P. Somnath and interns (A. Matthew, R. Bhave, J. Iyyadurai, A. Biswas, A. Manuel, P. Bramha, D. Manral, A. Sarkar, S. Totiger, H. Singh, A. Manohar, A. Kale, A. Jayanth, B. Pranav and K. Meenakshi) for help with data collection. We would also like to thank Department of Biotechnology – Indian Institute of Science (DBT-IISc) partnership program, Department of Science and Technology-Fund for Improvement of Science and Technology (DST-FIST) and the Science and Engineering Research Board (SERB Grant No. CRG/2018/003928) for funding this research, and the Ministry of Human Resource Development (MHRD) for providing fellowship during the course of study. The capturing and handling of animals used in the study was approved by the Institutional Animal Ethics Committee constituted by the Indian Institute of Science (CAF/ Ethics/390/2014).

## Supplementary material

**Table S1:** Results from mixed-effect models comparing female broadcast signals with that of males. Please see methods section for details of analysis. Females signal using head bobs more often while males use the relatively more conspicuous push ups and gular extension more than do females. Females are also seen in duller colour more commonly than are males, while males are observed in brighter colours more frequently. GLMMs with negative binomial error structure and log link were run for short duration behaviours (head bobs, push ups) while LMMs were run for long duration behaviours (gular extension, tail raise) and colour states. Estimates, 95% confidence intervals (CI) and results from likelihood ratio tests are shown. To increase confidence in the models, permutation tests for statistical null hypothesis testing of model coefficients were carried out with 1000 iterations. Additionally, model coefficients were bootstrapped with 1000 iterations.

|  | Head bobs |  |  |  |  |  |  |  | Push ups |  |  |  |  |  |  |  |
| --- | --- | --- | --- | --- | --- | --- | --- | --- | --- | --- | --- | --- | --- | --- | --- | --- |
| | Estimate | 95% CI | | $\chi^2$ | df | *p val* | Randomisa<br>tion *p val* | Bootstrapped<br>95% CI | Estimate | 95% CI | | $\chi^2$ | df | *p val* | Randomisa<br>tion *p val* | Bootstrapped<br>95% CI |
| Intercept<br>(Sex: Female) | -4.25 | -4.54 | -3.95 |  |  |  | 0.99 | -4.56 -4.02 | -7.52 | -8.22 | -6.82 |  |  |  | 0.03 | -8.38 -7.04 |
| Sex: Male | **-0.55** | **-0.98** | **-0.12** | **5.90** | **1** | **0.02** | **0.02** | **-1.24 -0.13** | **1.37** | **0.40** | **2.35** | **6.82** | **1** | **0.01** | **0.02** | **0.42 2.25** |
|  | Gular extension |  |  |  |  |  |  |  | Tail raise |  |  |  |  |  |  |  |
| | Estimate | 95% CI | | $\chi^2$ | df | *p val* | Randomisa<br>tion *p val* | Bootstrapped<br>95% CI | Estimate | 95% CI | | $\chi^2$ | df | *p val* | Randomisa<br>tion *p val* | Bootstrapped<br>95% CI |
| Intercept<br>(Sex: Female) | 0.01 | -0.04 | 0.05 |  |  |  | 1.00 | 0.003 0.01 | 0.20 | 0.14 | 0.26 |  |  |  | <0.01 | 0.12 0.28 |
| Sex: Male | **0.08** | **0.02** | **0.15** | **5.98** | **1** | **0.01** | **<0.01** | **0.02 0.16** | **-0.18** | **-0.27** | **-0.09** | **13.21** | **1** | **<0.01** | **<0.01** | **-0.27 -0.09** |
|  | Camouflaging colour |  |  |  |  |  |  |  | Conspicuous colour |  |  |  |  |  |  |  |
| | Estimate | 95% CI | | $\chi^2$ | df | *p val* | Randomisa<br>tion *p val* | Bootstrapped<br>95% CI | Estimate | 95% CI | | $\chi^2$ | df | *p val* | Randomisa<br>tion *p val* | Bootstrapped<br>95% CI |
| Intercept<br>(Sex: Female) | 0.90 | 0.78 | 1.03 |  |  |  | <0.01 | 0.83 0.97 | 0.09 | -0.04 | 0.22 |  |  |  | 1.00 | 0.04 0.16 |
| Sex: Male | **-0.56** | **-0.74** | **-0.38** | **27.35** | **1** | **<0.01** | **<0.01** | **-0.74 -0.37** | **0.50** | **0.31** | **0.68** | **22.50** | **1** | **<0.01** | **<0.01** | **0.33 0.68** |

**Table S2:**
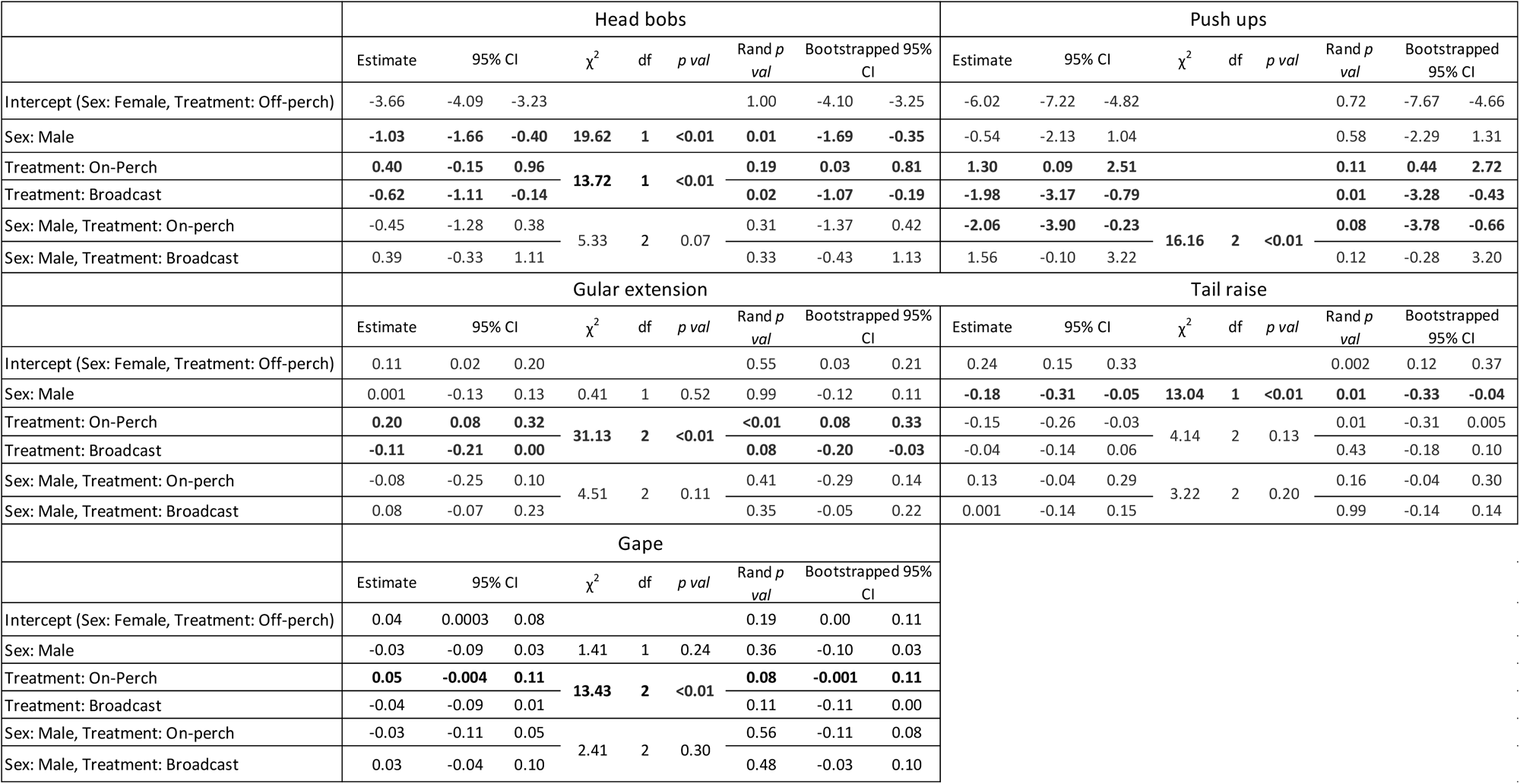
Results from mixed-effect models examining variation in signals shown by both males and females at varying levels of simulated intruder threat. (Please see analysis section for details of models used). Only females appear to increase frequency of head bobs and push ups while both sexes increase time spent in gular extension and gape with increasing levels of competitor threat. GLMMs with negative binomial error structure and log link were run for short duration behaviours (head bobs, push ups) and LMMs were run for long duration behaviours (gular extension, tail raise) and colour states. Estimates, 95% confidence intervals (CI) and results from likelihood ratio tests are shown. To increase confidence in the models, permutation tests for statistical null hypothesis testing of model coefficients were carried out with 1000 iterations. Additionally, model coefficients were bootstrapped with 1000 iterations. When interactions were statistically significant, the statistical significance of main effects was not reported since they are not meaningful in our study.

**Table S3:** Results from linear mixed effects models (LMMs) examining variation in proportion of time spent in colour and behaviour states at varying levels of simulated intruder threat unique to one sex. Females do not appear to vary time spent in different colour states with varying level of competitor threat. In contrast, males increased duration in dorsal yellow-lateral red colour and a laterally compressed posture. Estimates, 95 % confidence intervals, statistics from likelihood ratio tests are shown. To increase confidence in the models, permutation tests for assessment of p values were carried out and coefficients were bootstrapped.

|  | Grey body-orange head (females) |  |  |  |  |  |  | Dark body-orange head (females) |  |  |  |  |  |  |
| --- | --- | --- | --- | --- | --- | --- | --- | --- | --- | --- | --- | --- | --- | --- |
| | Estimate | 95% CI | $\chi^2$ | df | p val | Randomisat<br>ion p val | Bootstrapped<br>95% CI | Estimate | 95% CI | $\chi^2$ | df | p val | Randomisa<br>tion p val | Bootstrapped<br>95% CI |
| Intercept (Treatment:<br>Off-perch) | 0.30 | 0.13 0.47 |  |  |  | 0.78 | 0.16 0.48 | 0.30 | 0.16 0.44 |  |  |  | 0.04 | 0.16 0.47 |
| Treatment: On-Perch | 0.05 | -0.15 0.24 |  |  |  | 0.67 | -0.14 0.25 | -0.11 | -0.28 0.06 |  |  |  | 0.21 | -0.33 0.11 |
| Treatment: Broadcast | 0.07 | -0.10 0.24 | 0.63 | 2 | 0.73 | 0.42 | -0.06 0.20 | -0.14 | -0.29 0.01 | 3.49 | 2 | 0.17 | 0.06 | -0.28 -0.02 |
|  | Dorsal yellow colour (males) |  |  |  |  |  |  | Dorsal orange colour (males) |  |  |  |  |  |  |
| | Estimate | 95% CI | $\chi^2$ | df | p val | Randomisat<br>ion p val | Bootstrapped<br>95% CI | Estimate | 95% CI | $\chi^2$ | df | p val | Randomisa<br>tion p val | Bootstrapped<br>95% CI |
| Intercept (Treatment:<br>Off-perch) | 0.46 | 0.30 0.63 |  |  |  | 0.36 | 0.30 0.63 | 0.10 | -0.02 0.22 |  |  |  | 0.88 | 0.002 0.23 |
| Treatment: On-Perch | -0.02 | -0.19 0.15 |  |  |  | 0.85 | -0.18 0.15 | 0.09 | -0.04 0.23 |  |  |  | 0.17 | -0.07 0.26 |
| Treatment: Broadcast | -0.03 | -0.17 0.12 | 0.12 | 2 | 0.94 | 0.74 | -0.17 0.10 | 0.05 | -0.06 0.17 | 1.99 | 2 | 0.37 | 0.39 | -0.03 0.15 |
|  | Dorsal yellow-lateral red colour (males) |  |  |  |  |  |  | Lateral compression (males) |  |  |  |  |  |  |
| | Estimate | 95% CI | $\chi^2$ | df | p val | Randomisat<br>ion p val | Bootstrapped<br>95% CI | Estimate | 95% CI | $\chi^2$ | df | p val | Randomisa<br>tion p val | Bootstrapped<br>95% CI |
| Intercept (Treatment:<br>Off-perch) | 0.02 | -0.03 0.07 |  |  |  | 0.65 | -5.90 0.05 | 0.23 | 0.08 0.37 |  |  |  | 0.53 | 0.09 0.38 |
| Treatment: On-Perch | **0.09** | **0.01 0.17** |  |  |  | **0.03** | **-0.01 0.20** | **0.16** | **-0.02 0.33** |  |  |  | **0.07** | **-0.08 0.38** |
| Treatment: Broadcast | -0.02 | -0.08 0.05 | 9.59 | 2 | 0.01 | 0.65 | -0.05 0.004 | -0.06 | -0.21 0.08 | 8.11 | 2 | 0.02 | 0.40 | -0.18 0.03 |

### Control experiments

#### Data collection (control model presentation)

A control experiment was performed to control for lizard response to a novel object on their erritory (i.e., to check whether lizard responses to the lizard models represented responses to onspecifics or to a novel object on their territory). This experiment also allowed us to examine whether lizard responses were in response to observer movement towards the focal individual while placing the model at the desired location. A grey PVC pipe (Figure S3) was presented as a ontrol to 14 females between 2015 and 2017 and to 13 males between 2013 and 2016. The control model was presented to the focal individual on the perch of the focal individual. Female model presentation was carried out on the typically used perch identified by observers while male presentation was carried out at the perch the focal male was using 5-10 minutes prior to starting the ocal observation. Every individual presented with a control model on perch was presented with a izard model on perch for comparison. Care was taken that a control model and lizard model were presented within 10 days of each other, and the sequence of presentation was randomised. Control model presentation was preceded by a 30 minute focal in the natural setting, without manipulating ts environment (broadcast signalling) and the procedure of model presentation and recording of behaviours was the same as described for the intruder model presentation in the main manuscript. Videos were transcribed using software BORIS and rates of body movements, and proportion of ime spent in postures and colours were compared between natural observations, on-perch control model presentation and on-perch lizard model presentations. Responses of males and females were ompared separately.

#### Statistical analysis (control model presentation)

Generalised linear mixed effects models with negative binomial error structure, duration of focal as n offset term and identity of the individual as a random effect were run for females and males eparately with counts of head bobs and push ups as the response variables and treatment type Broadcast signalling, control model presentation, lizard model on perch) as predictor. LMEs with imilar fixed and random effects were run for posture and colour signals, which are quantified using proportion of time spent. Comparison between control model and lizard model presentation helped understand whether individuals perceived lizard models as competitors or as any other novel object in their territory. Comparison between observation when no model was presented, and ontrol model presentation helped understand if the whole process of an observer walking towards he focal individual to present the model affected the behaviour of individuals. Permutation tests were run and 95% bootstrapped confidence intervals were calculated to increase our confidence in models.

#### Results (control model presentation)

Comparing signalling between presentations of on-perch lizard models and on-perch control models in both females and males revealed that females substantially increase use of head bobs, push ups, gular extension and gape while males increased use of gular extension, gape, lateral ompression and yellow-red colouration when presented with a lizard model as compared to a ontrol model (Figure S1, S2, Table S4, S5).

### 1. Females

**Figure S1:**
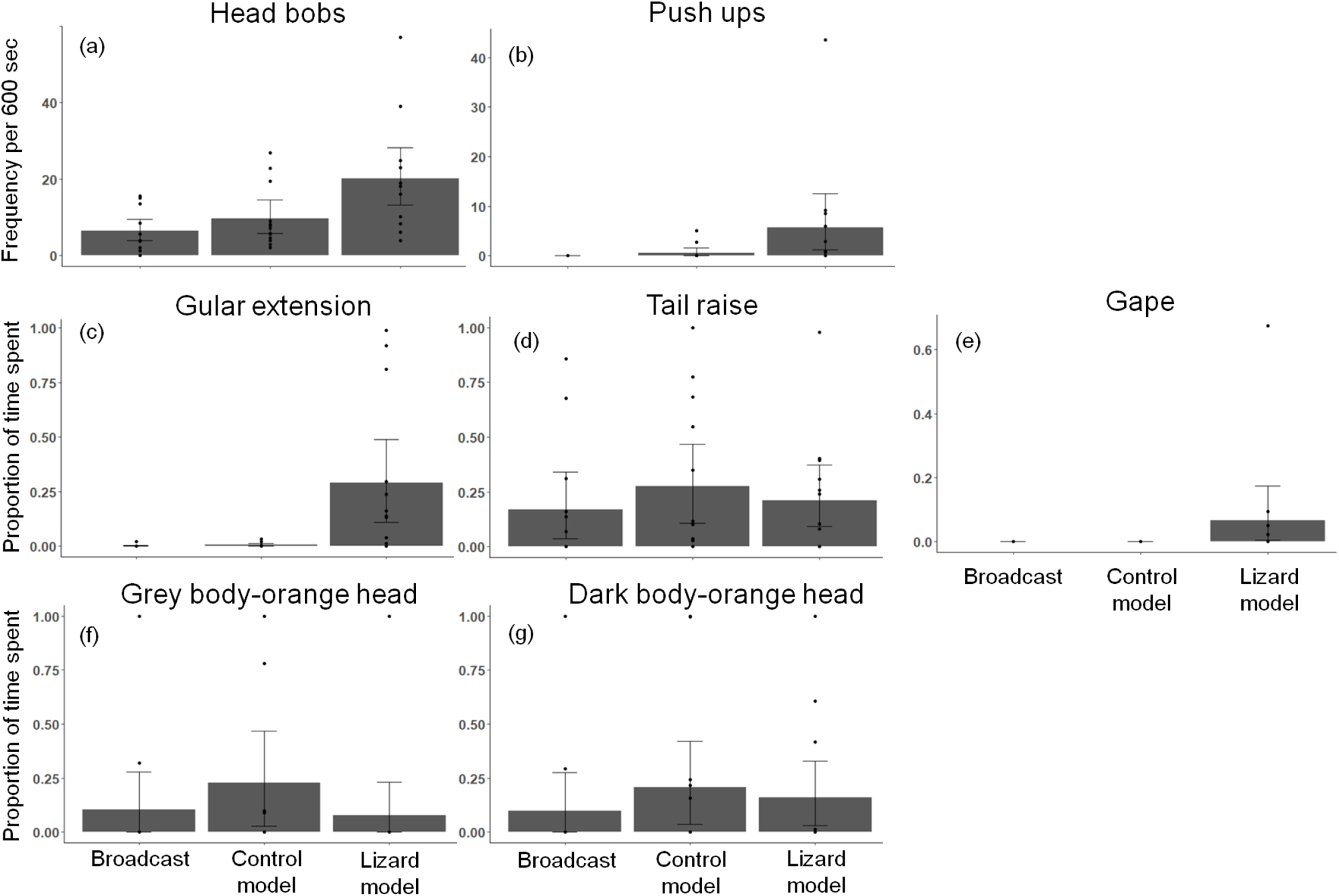
Comparison between broadcast signalling, signalling on simulation of a control model and a lizard model in females. Females appear to substantially increase frequency of head bobs (a) and push ups (b) and duration of gular extension (c) and gape (e) when presented with a lizard model as compared to a control model. Signalling rates and time spent in postures and colours did not appear to vary during control model simulation and broadcast signalling (a-g). Error bars represent 95% bootstrapped confidence intervals, while the dots represent the raw data points

**Table S4:**
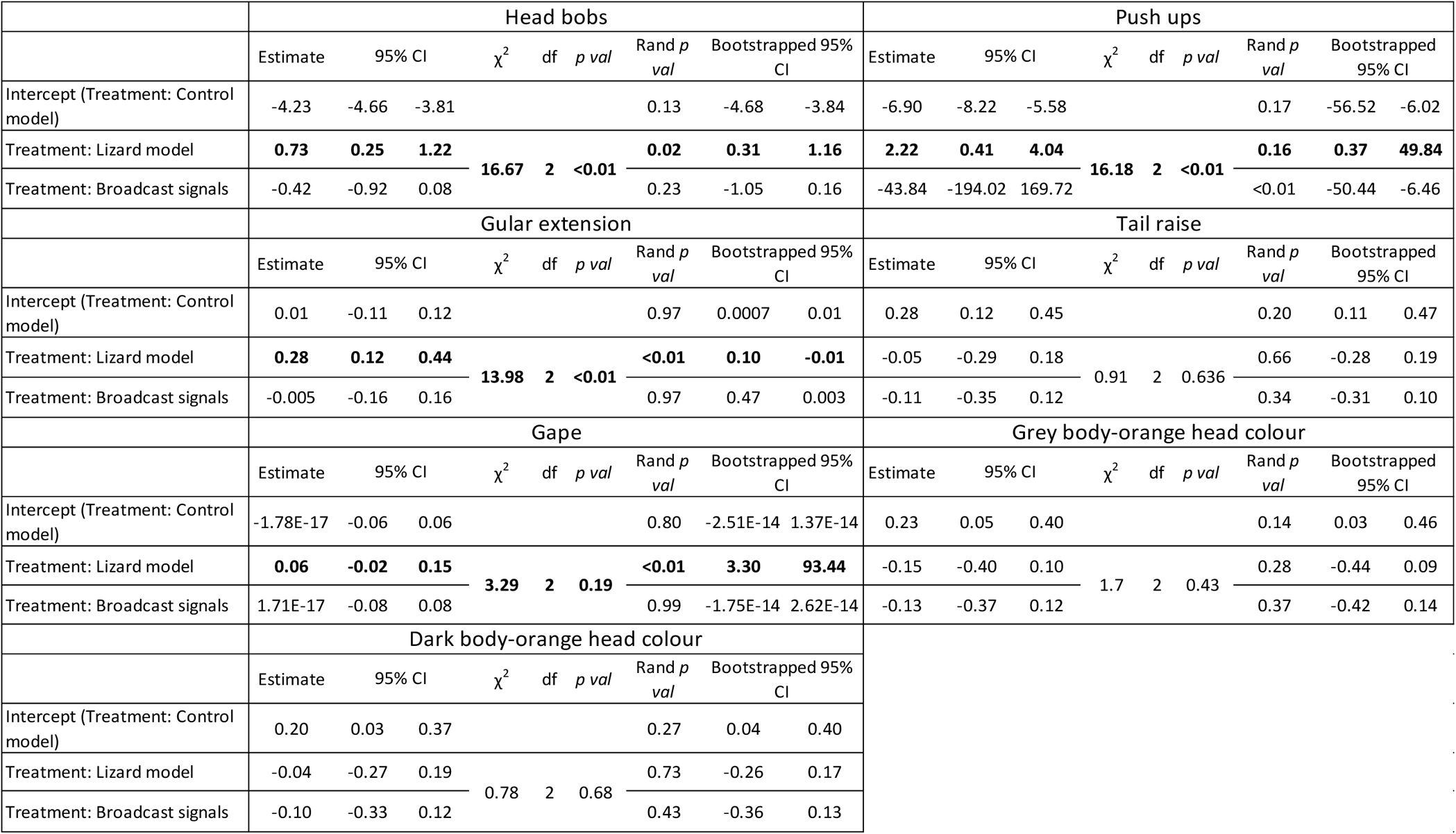
Results from models examining variation in frequencies (head bobs, push ups) and proportion of time spent (gular extension, tail raise, gape) on simulation of an on-perch control model, on-perch lizard model and broadcast signalling in *females* (when no model was presented). Females appear to increase frequency of head bobs and push ups and duration of gular extension and gape when presented with a lizard model as compared to a control model. No statistically significant difference in signalling was observed between control model and broadcast signals. GLMMs with negative binomial error structure and log link were run for short duration behaviours (head bobs, push ups) while LMMs were run for long duration behaviours (gular extension, tail raise, gape) and colour states. Estimates, 95% confidence intervals (CI) and results from likelihood ratio tests are shown. To increase confidence in the models, permutation tests for statistical null hypothesis testing of model coefficients were carried out with 1000 iterations. Additionally, model coefficients were bootstrapped with 1000 iterations.

|  | Head bobs |  |  |  |  |  |  |  | Push ups |  |  |  |  |  |  |  |
| --- | --- | --- | --- | --- | --- | --- | --- | --- | --- | --- | --- | --- | --- | --- | --- | --- |
| | Estimate | 95% CI | | $\chi^2$ | df | p val | Rand p val | Bootstrapped 95% CI | Estimate | 95% CI | | $\chi^2$ | df | p val | Rand p val | Bootstrapped 95% CI |
| Intercept (Treatment: Control model) | -4.23 | -4.66 | -3.81 |  |  |  | 0.13 | -4.68 -3.84 | -6.90 | -8.22 | -5.58 |  |  |  | 0.17 | -56.52 -6.02 |
| Treatment: Lizard model | **0.73** | **0.25** | **1.22** | **16.67** | **2** | **<0.01** | **0.02** | **0.31 1.16** | **2.22** | **0.41** | **4.04** | **16.18** | **2** | **<0.01** | **0.16** | **0.37 49.84** |
| Treatment: Broadcast signals | -0.42 | -0.92 | 0.08 |  |  |  | 0.23 | -1.05 0.16 | -43.84 | -194.02 | 169.72 |  |  |  | <0.01 | -50.44 -6.46 |
|  | Gular extension |  |  |  |  |  |  |  | Tail raise |  |  |  |  |  |  |  |
| | Estimate | 95% CI | | $\chi^2$ | df | p val | Rand p val | Bootstrapped 95% CI | Estimate | 95% CI | | $\chi^2$ | df | p val | Rand p val | Bootstrapped 95% CI |
| Intercept (Treatment: Control model) | 0.01 | -0.11 | 0.12 |  |  |  | 0.97 | 0.0007 0.01 | 0.28 | 0.12 | 0.45 |  |  |  | 0.20 | 0.11 0.47 |
| Treatment: Lizard model | **0.28** | **0.12** | **0.44** | **13.98** | **2** | **<0.01** | **<0.01** | **0.10 -0.01** | -0.05 | -0.29 | 0.18 | 0.91 | 2 | 0.636 | 0.66 | -0.28 0.19 |
| Treatment: Broadcast signals | -0.005 | -0.16 | 0.16 |  |  |  | 0.97 | 0.47 0.003 | -0.11 | -0.35 | 0.12 |  |  |  | 0.34 | -0.31 0.10 |
|  | Gape |  |  |  |  |  |  |  | Grey body-orange head colour |  |  |  |  |  |  |  |
| | Estimate | 95% CI | | $\chi^2$ | df | p val | Rand p val | Bootstrapped 95% CI | Estimate | 95% CI | | $\chi^2$ | df | p val | Rand p val | Bootstrapped 95% CI |
| Intercept (Treatment: Control model) | -1.78E-17 | -0.06 | 0.06 |  |  |  | 0.80 | -2.51E-14 1.37E-14 | 0.23 | 0.05 | 0.40 |  |  |  | 0.14 | 0.03 0.46 |
| Treatment: Lizard model | **0.06** | **-0.02** | **0.15** | **3.29** | **2** | **0.19** | **<0.01** | **3.30 93.44** | -0.15 | -0.40 | 0.10 | 1.7 | 2 | 0.43 | 0.28 | -0.44 0.09 |
| Treatment: Broadcast signals | 1.71E-17 | -0.08 | 0.08 |  |  |  | 0.99 | -1.75E-14 2.62E-14 | -0.13 | -0.37 | 0.12 |  |  |  | 0.37 | -0.42 0.14 |
|  | Dark body-orange head colour |  |  |  |  |  |  |  |  |  |  |  |  |  |  |  |
| | Estimate | 95% CI | | $\chi^2$ | df | p val | Rand p val | Bootstrapped 95% CI | | | | | | | | |
| Intercept (Treatment: Control model) | 0.20 | 0.03 | 0.37 |  |  |  | 0.27 | 0.04 0.40 |  |  |  |  |  |  |  |  |
| Treatment: Lizard model | -0.04 | -0.27 | 0.19 | 0.78 | 2 | 0.68 | 0.73 | -0.26 0.17 |  |  |  |  |  |  |  |  |
| Treatment: Broadcast signals | -0.10 | -0.33 | 0.12 |  |  |  | 0.43 | -0.36 0.13 |  |  |  |  |  |  |  |  |

### 2. Males

**Figure S2:**
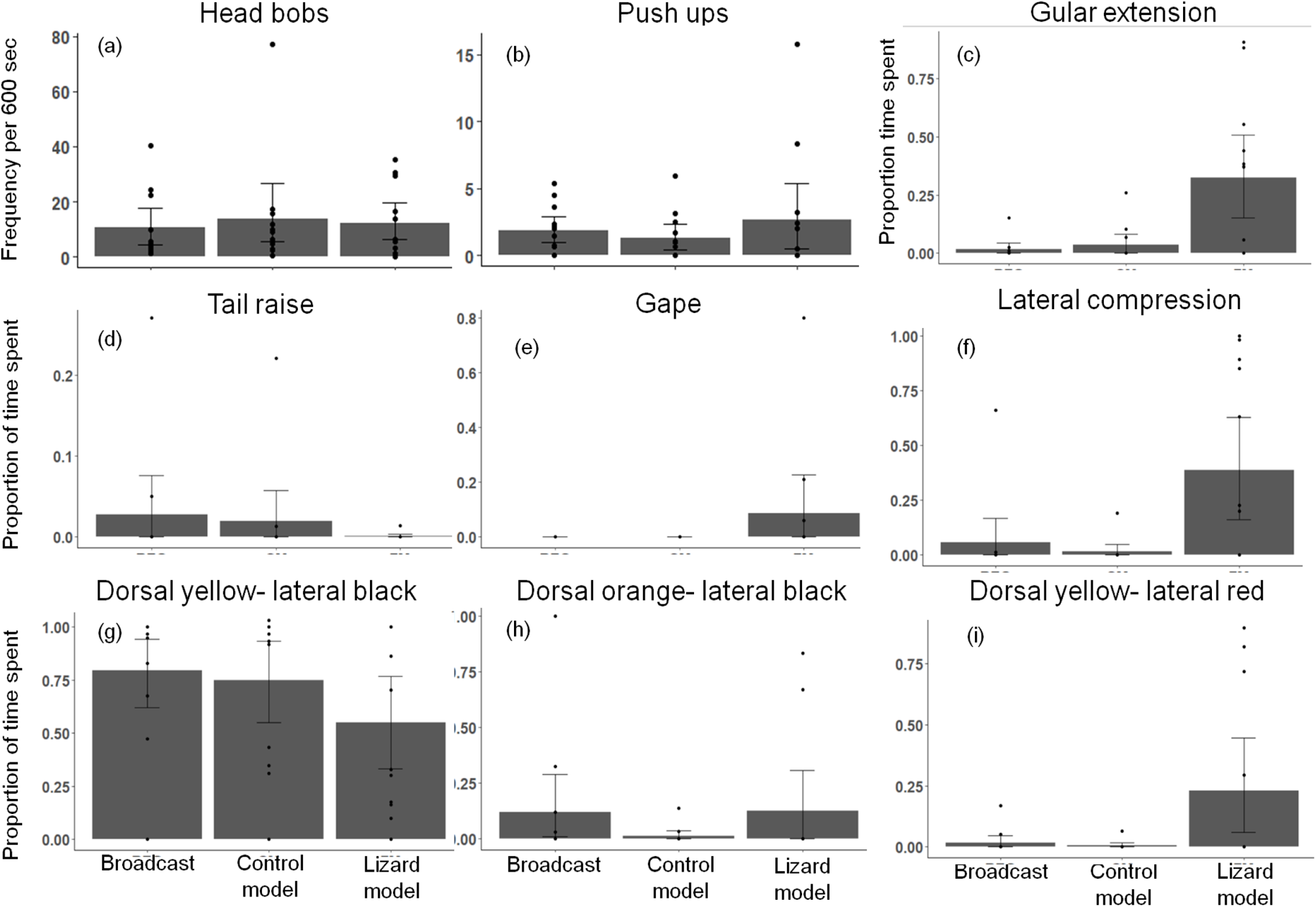
Comparison between male signalling on simulation of an on-perch control model, on-perch lizard model and broadcast signalling (when no model was presented). Males appear to substantially increase time spent in gular extension (c), gape (e), lateral compression (f) and in dorsal yellow-lateral red colour (g) when presented with a lizard model as compared to a control model. No statistically significant difference in signalling was observed between control model and broadcast signals (a-i). Error bars represent 95% bootstrapped confidence intervals, while the dots represent the actual data points

**Table S5:**
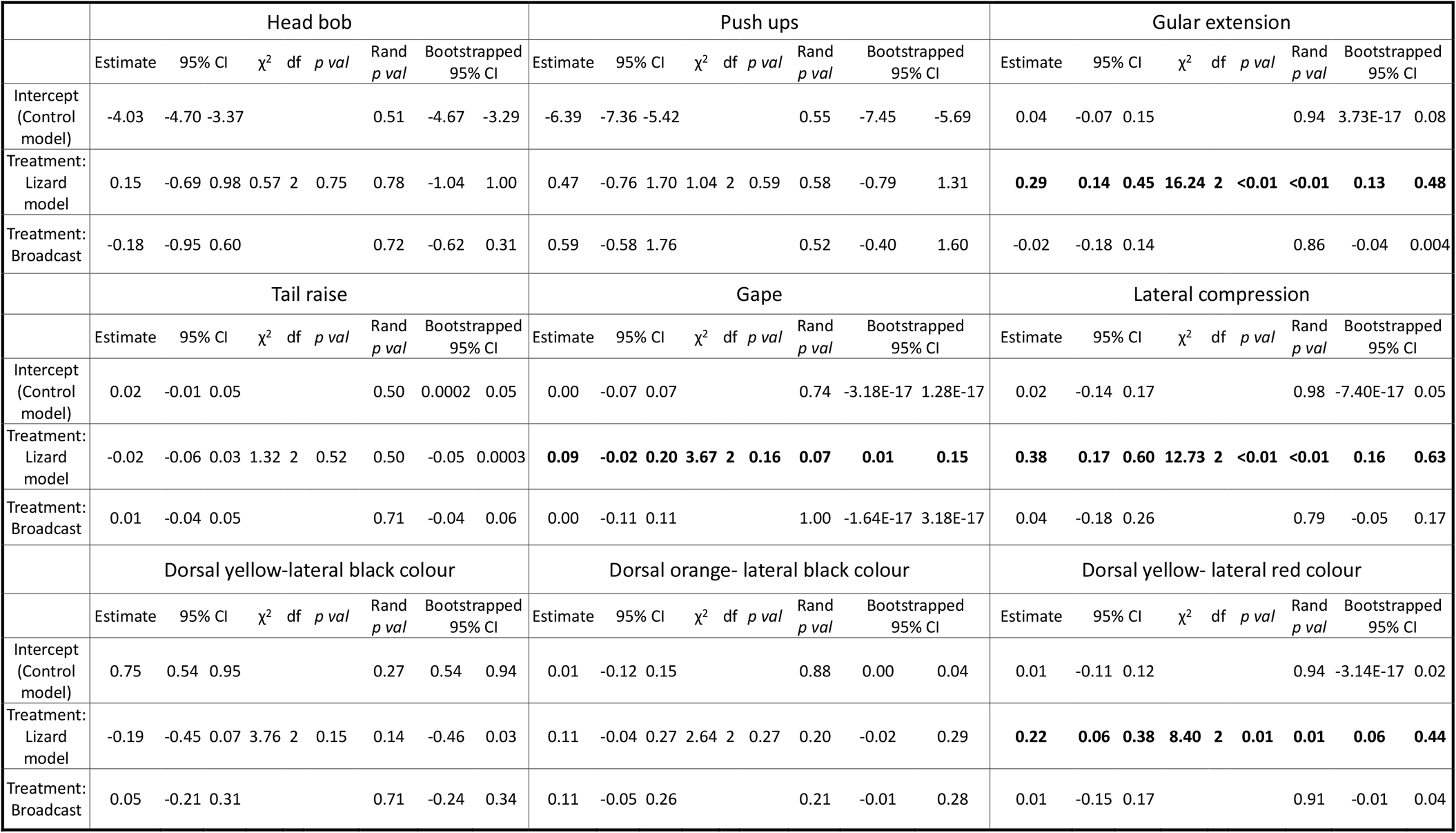
Results from models examining variation in frequencies (head bobs, push ups) and proportion of time spent (gular extension, tail raise, gape) on simulation of an on-perch control model, on-perch lizard model and broadcast signalling in *males* (when no model was presented). Males appear to increase time spent in gular extension, gape, lateral compression and in dorsal yellow-lateral red colour when presented with a lizard model as compared to a control model. No statistically significant difference in signalling was observed between control model and broadcast signals. GLMMs with negative binomial error structure and log link were run for short duration behaviours (head bobs, push ups) while LMMs were run for long duration behaviours (gular extension, tail raise, gape, lateral compression) and colour states. Estimates, 95% confidence intervals (CI) and results from likelihood ratio tests are shown. To increase confidence in the models, permutation tests for statistical null hypothesis testing of model coefficients were carried out with 1000 iterations. Additionally, model coefficients were bootstrapped with 1000 iterations.

|  | Head bob |  |  |  |  |  |  |  | Push ups |  |  |  |  |  |  |  | Gular extension |  |  |  |  |  |  |  |
| --- | --- | --- | --- | --- | --- | --- | --- | --- | --- | --- | --- | --- | --- | --- | --- | --- | --- | --- | --- | --- | --- | --- | --- | --- |
| | Estimate | 95% CI | $\chi^2$ | df | p val | Rand p val | Bootstrapped 95% CI | | Estimate | 95% CI | $\chi^2$ | df | p val | Rand p val | Bootstrapped 95% CI | | Estimate | 95% CI | $\chi^2$ | df | p val | Rand p val | Bootstrapped 95% CI | |
| Intercept (Control model) | -4.03 | -4.70 -3.37 |  |  |  | 0.51 | -4.67 -3.29 |  | -6.39 | -7.36 -5.42 |  |  |  | 0.55 | -7.45 -5.69 |  | 0.04 | -0.07 0.15 |  |  |  | 0.94 | 3.73E-17 | 0.08 |
| Treatment: Lizard model | 0.15 | -0.69 0.98 | 0.57 | 2 | 0.75 | 0.78 | -1.04 1.00 |  | 0.47 | -0.76 1.70 | 1.04 | 2 | 0.59 | 0.58 | -0.79 1.31 |  | **0.29** | **0.14 0.45** | **16.24** | **2** | **<0.01** | **<0.01** | **0.13** | **0.48** |
| Treatment: Broadcast | -0.18 | -0.95 0.60 |  |  |  | 0.72 | -0.62 0.31 |  | 0.59 | -0.58 1.76 |  |  |  | 0.52 | -0.40 1.60 |  | -0.02 | -0.18 0.14 |  |  |  | 0.86 | -0.04 | 0.004 |
|  | Tail raise |  |  |  |  |  |  |  | Gape |  |  |  |  |  |  |  | Lateral compression |  |  |  |  |  |  |  |
| | Estimate | 95% CI | $\chi^2$ | df | p val | Rand p val | Bootstrapped 95% CI | | Estimate | 95% CI | $\chi^2$ | df | p val | Rand p val | Bootstrapped 95% CI | | Estimate | 95% CI | $\chi^2$ | df | p val | Rand p val | Bootstrapped 95% CI | |
| Intercept (Control model) | 0.02 | -0.01 0.05 |  |  |  | 0.50 | 0.0002 0.05 |  | 0.00 | -0.07 0.07 |  |  |  | 0.74 | -3.18E-17 1.28E-17 |  | 0.02 | -0.14 0.17 |  |  |  | 0.98 | -7.40E-17 | 0.05 |
| Treatment: Lizard model | -0.02 | -0.06 0.03 | 1.32 | 2 | 0.52 | 0.50 | -0.05 0.0003 |  | **0.09** | **-0.02 0.20** | **3.67** | **2** | **0.16** | **0.07** | **0.01 0.15** |  | **0.38** | **0.17 0.60** | **12.73** | **2** | **<0.01** | **<0.01** | **0.16** | **0.63** |
| Treatment: Broadcast | 0.01 | -0.04 0.05 |  |  |  | 0.71 | -0.04 0.06 |  | 0.00 | -0.11 0.11 |  |  |  | 1.00 | -1.64E-17 3.18E-17 |  | 0.04 | -0.18 0.26 |  |  |  | 0.79 | -0.05 | 0.17 |
|  | Dorsal yellow-lateral black colour |  |  |  |  |  |  |  | Dorsal orange- lateral black colour |  |  |  |  |  |  |  | Dorsal yellow- lateral red colour |  |  |  |  |  |  |  |
| | Estimate | 95% CI | $\chi^2$ | df | p val | Rand p val | Bootstrapped 95% CI | | Estimate | 95% CI | $\chi^2$ | df | p val | Rand p val | Bootstrapped 95% CI | | Estimate | 95% CI | $\chi^2$ | df | p val | Rand p val | Bootstrapped 95% CI | |
| Intercept (Control model) | 0.75 | 0.54 0.95 |  |  |  | 0.27 | 0.54 0.94 |  | 0.01 | -0.12 0.15 |  |  |  | 0.88 | 0.00 0.04 |  | 0.01 | -0.11 0.12 |  |  |  | 0.94 | -3.14E-17 | 0.02 |
| Treatment: Lizard model | -0.19 | -0.45 0.07 | 3.76 | 2 | 0.15 | 0.14 | -0.46 0.03 |  | 0.11 | -0.04 0.27 | 2.64 | 2 | 0.27 | 0.20 | -0.02 0.29 |  | **0.22** | **0.06 0.38** | **8.40** | **2** | **0.01** | **0.01** | **0.06** | **0.44** |
| Treatment: Broadcast | 0.05 | -0.21 0.31 |  |  |  | 0.71 | -0.24 0.34 |  | 0.11 | -0.05 0.26 |  |  |  | 0.21 | -0.01 0.28 |  | 0.01 | -0.15 0.17 |  |  |  | 0.91 | -0.01 | 0.04 |

#### The models

Lizard models used were 3D printed, painted using acrylic colours and stationary (Supplementary Figure S3). Stationary models allowed us to simulate a similar morphological and behavioural phenotype to all focal individuals, which tethering of live individuals or presenting live individuals in a cage would not have allowed. To control for size effects, the snout-vent length of models was size matched (+/− 1 cm) to the snout-vent length of the focal individuals. Because *P. dorsalis* exhibits dynamic colour change, we had to decide which colour to use for the models. We used the colour that was dominant during the mating season. Females spent 74% of time in grey colour with an off-white head (this study) while males spent 45% of their time in yellow colour dorsally and black laterally (males spent 30% of their time in pale yellow-light black colour and 15% of their time in dorsal orange-lateral black colour, unpublished data) during the mating season. Thus, models presented to females were grey in colour (Supplementary Figure S3a) while those presented to males were painted yellow dorsally and black laterally (Supplementary Figure S3b).

**Figure S3:**
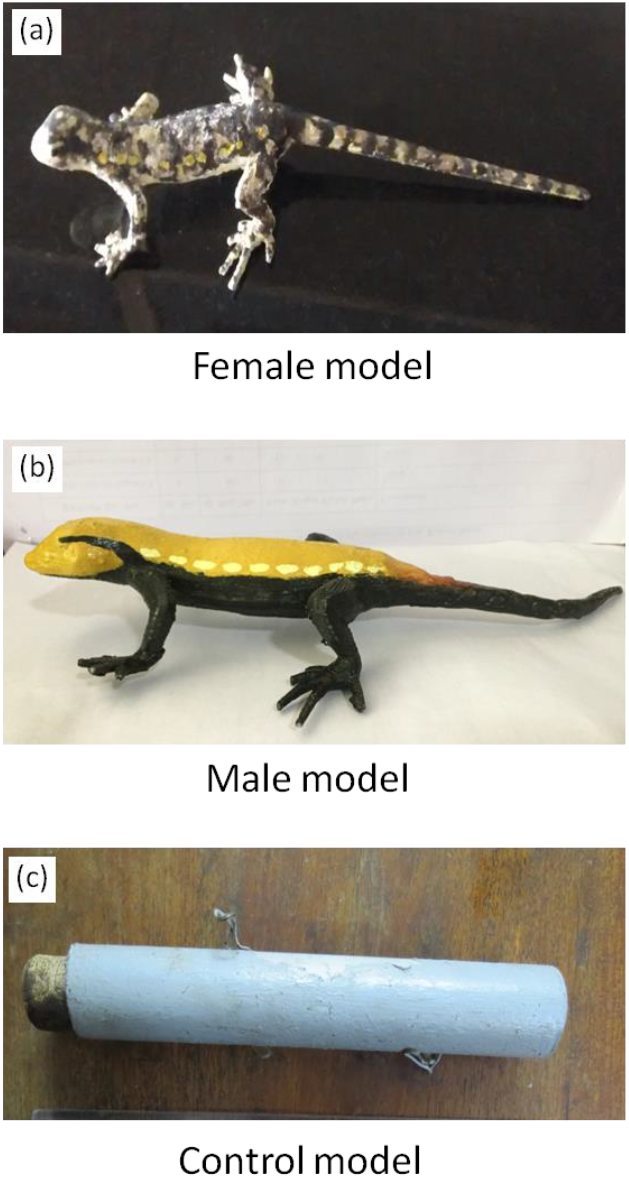
Models used in simulating intruder threat to females (a) and males (b) and in control experiments on both sexes (c)

## References

Altmann, J. (1974). Observational Study of Behavior: Sampling Methods. Source Behav., 49, 227–266.

Archie, E.A., Morrison, T.A., Foley, C.A.H., Moss, C.J. & Alberts, S.C. (2006). Dominance rank relationships among wild female African elephants, Loxodonta africana. Anim. Behav., 71, 117–127.

Baird, T. A. & Sloan, C. L. (2003). Interpopulation variation in the social organization of female collared lizards, *Crotaphytus collaris*. Ethology, 109, 879–894.

Baniel, A., Cowlishaw, G. & Huchard, E. (2018). Jealous females? Female competition and reproductive suppression in a wild promiscuous primate. Proc. R. Soc. B Biol. Sci., 285.

Batabyal, A. & Thaker, M. (2017). Signalling with physiological colours: high contrast for courtship but speed for competition. Anim. Behav., 129, 229–236.

Bebié, N. & McElligott, A. G. (2006). Female aggression in red deer: Does it indicate competition for mates? Mamm. Biol., 71, 347–355.

Bhadra, A., Mitra, A., Deshpande, S. A., Chandrasekhar, K., Naik, D. G., Hefetz, A., et al. (2010). Regulation of reproduction in the primitively eusocial wasp *Ropalidia marginata*: on the trail of the queen pheromone. J. Chem. Ecol., 36, 424–31.

Brooks, M. E., Kristensen, K., van Benthem, K. J., Magnusson, A., Berg, C. W., Nielsen, A., et al. (2017). Modeling Zero-Inflated Count Data With glmmTMB. bioRxiv, 132753.

Cant, M. A & Young, A. J. (2013). Resolving social conflict among females without overt aggression. Philos. Trans. R. Soc. Lond. B. Biol. Sci., 368, 20130076.

Chandrashekara, K. & Gadagkar, R. (1992). Queen succession in the primitively eusocial tropical wasp *Ropalidia marginata* (Lep.) (Hymenoptera: Vespidae). J. Insect Behav., 5, 193–209.

Clutton-Brock, T.H. (2007). Sexual selection in males and females. Science (80-.)., 318, 1882–1885.

Clutton-Brock, T.H. (2009). Sexual selection in females. Anim. Behav., 77, 3–11.

Clutton-Brock, T. H., Hodge, S. J., Spong, G., Russell, A. F., Jordan, N. R., Bennett, N. C., et al. (2006). Intrasexual competition and sexual selection in cooperative mammals. Nature, 444, 1065–1068.

Clutton-Brock, T.H. & Huchard, E. (2013a). Social competition and its consequences in female mammals. J. Zool., 289, 151–171.

Clutton-Brock, T.H. & Huchard, E. (2013b). Social competition and selection in males and females. Philos. Trans. R. Soc. Lond. B. Biol. Sci., 368, 20130074.

Daniel, J. C. (2002). The Book of Indian Reptiles and Amphibians. Bombay Natural History Society and Oxford University Press.

Darwin, C. (1871). The descent of man and selection in relation to sex. 1st edn. John Murray, London.

Deodhar, S. & Isvaran, K. (2017). Breeding phenology of Psammophilus dorsalis : patterns in time, space and morphology. Curr. Sci., 113, 2120–2126.

Deodhar, S. & Isvaran, K. (2018). Why Do Males Use Multiple Signals? Insights From Measuring Wild Male Behavior Over Lifespans. Front. Ecol. Evol., 6, 1–12.

Diniz, P., Oliveira, R.S., Marini, M. & Duca, C. (2019). Angry caciques: intrasexual aggression in a Neotropical colonial blackbird. Ethol. Ecol. Evol., 31, 205–218.

Draud, M., Macías-Ordóñez, R., Verga, J. & Itzkowitz, M. (2004). Female and male Texas cichlids *(Herichthys cyanoguttatum)* do not fight by the same rules. Behav. Ecol., 15, 102–108.

Eggert, A., Otte, T. & Müller, J. (2008). Starving the competition: a proximate cause of reproductive skew in burying beetles (Nicrophorus vespilloides). Proc. R. Soc. B.

Elias, D. O., Botero, C. A., Andrade, M. C. B., Mason, A. C. & Kasumovic, M. M. (2010). High resource valuation fuels “desperado” fighting tactics in female jumping spiders. Behav. Ecol., 21, 868–875.

Friard, O. & Gamba, M. (2016). BORIS: a free, versatile open-source event-logging software for video/audio coding and live observations. Methods Ecol. Evol.

Geberzahn, N., Goymann, W., Muck, C. & ten Cate, C. (2009). Females alter their song when challenged in a sex-role reversed bird species. Behav. Ecol. Sociobiol., 64, 193–204.

Haenel, G. J., Smith, L. C. & John-Alder, H. B. (2003). Home-Range Analysis in *Sceloporus undulatus* (Eastern Fence Lizard). I. Spacing Patterns and the Context of Territorial Behavior. Copeia, 2003, 99–112.

Hare, R. M. & Simmons, L. W. (2019). Sexual selection and its evolutionary consequences in female animals. Biol. Rev., 94, 929–956.

Hodge, S. J., Flower, T. P. & Clutton-Brock, T. H. (2007). Offspring competition and helper associations in cooperative meerkats. Anim. Behav., 74, 957–964.

Hughes, M. (1996). Size assessment via a visual signal in snapping shrimp. Behav. Ecol. Sociobiol., 38, 51–57.

Johnson, K. (1988). Sexual selection in pinyon jays II: male choice and female-female competition. Anim. Behav., 36, 1048–1053.

Kokita, T. (2002). The role of female behavior in maintaining monogamy of a coral-reef filefish. Ethology, 108, 157–168.

Krieg, C. A. & Getty, T. (2020). Fitness benefits to intrasexual aggression in female house wrens, *Troglodytes aedon*. Anim. Behav., 160, 79–90.

Kutsukake, N. & Clutton-Brock, T. H. (2006). Aggression and submission reflect reproductive conflict between females in cooperatively breeding meerkats *Suricata suricatta*. Behav. Ecol. Sociobiol., 59, 541–548.

Moran, M. (2003). Arguments for Rejecting the Sequential Bonferroni in Ecological Studies. Oikos, 100, 403–405.

Papadopoulos, N. T., Carey, J. R., Liedo, P., Müller, H. G. & Sentürk, D. (2009). Virgin females compete for mates in the male lekking species *Ceratitis capitata*. Physiol. Entomol., 34, 238–245.

R Core Team. (2014). R: A Language and Environment for Statistical Computing. R Foundation for Statistical Computing, Vienna, Austria.

Radder, R. S., Saidapur, S. K. & Shanbhag, B. A. (2006). Big boys on top: effects of body size, sex and reproductive state on perching behaviour in the tropical rock dragon, *Psammophilus dorsalis*. Anim. Biol., 56, 311–321.

Ranade, D. & Isvaran, K. (2022). Inferring Social Interactions Over a Lifespan from Space-Use Patterns in a Tropical Agamid. J. Herpetol., 56, 164–171.

Ratnieks, F. L.W., Foster, K. R. & Wenseleers, T. (2006). Conflict Resolution in Insect Societies. Annu. Rev. Entomol., 51, 581–608.

Reece, S. E., Innocent, T. M. & West, S. A. (2007). Lethal male-male combat in the parasitoid *Melittobia acasta*: are size and competitive environment important? Anim. Behav., 74, 1163–1169.

Robles, C. & Halloy, M. (2009). Home Ranges and Reproductive Strategies in a Neotropical Lizard, Liolaemus quilmes (Iguania: Liolaemidae). South Am. J. Herpetol., 4, 253–258.

Rosvall, K.A. (2011). Intrasexual competition in females: evidence for sexual selection? Behav. Ecol., 22, 1131–1140.

Schofield, G., Katselidis, K.A., Pantis, J.D., Dimopoulos, P. & Hays, G.C. (2007). Female-female aggression in loggerhead seaturtles. Mar. Ecol. Prog. Ser., 336, 267–274.

Scott, E. M., Mann, J., Watson-Capps, J. J., Sargeant, B. L. & Connor, R. C. (2005). Aggression in bottlenose dolphins: Evidence for sexual coercion, male-male competition, and female tolerance through analysis of tooth-rake marks and behaviour. Behaviour, 142, 21–44.

Shelly, T. (1999). Defense of Oviposition Sites by Female Oriental Fruit Flies (Diptera: Tephritidae). Florida Entomol., 82, 339–346.

Shuster, S.M. & Wade, M.J. (2003). Mating systems and strategies. Princeton University Press.

Smith, M. A. (1935). The Fauna of British India, including Ceylon and Burma. Taylor And Francis, Red Lion Court; London, 1935.

Stockley, P. & Bro-Jørgensen, J. (2011). Female competition and its evolutionary consequences in mammals. Biol. Rev., 86, 341–366.

Stockley, P. & Campbell, A. (2013). Female competition and aggression: interdisciplinary perspectives. Philos. Trans. R. Soc. Lond. B. Biol. Sci., 368, 20130073.

Stuart-smith, J., Swain, R. & Wapstral, E. (2007). The Role of Body Size in Competition and Mate Choice in an Agamid with Female-Biased Size Dimorphism. Behaviour, 144, 1087–1102.

Summers, K. (1989). Sexual selection and intra-female competition in the green poison-dart frog, *Dendrobates auratus*. Anim. Behav., 37, 797–805.

Tobias, J.A., Montgomerie, R. & Lyon, B.E. (2012). The evolution of female ornaments and weaponry: social selection, sexual selection and ecological competition. Philos. Trans. R. Soc. B Biol. Sci., 367, 2274–2293.

West-Eberhard, M.J. (1979). Sexual selection, social competition, and evolution. Evolution (N. Y)., 123, 222–234.

West-Eberhard, M.J. (2014). Darwin’s forgotten idea: The social essence of sexual selection. Neurosci. Biobehav. Rev., 46, 501–508.

While, G. M., Sinn, D. L. & Wapstra, E. (2009). Female aggression predicts mode of paternity acquisition in a social lizard. Proc. R. Soc. London B Biol. Sci., 276, 2021–2029.

Young, A. J., Carlson, A. A., Monfort, S. L., Russell, A. F., Bennett, N. C. & Clutton-Brock, T. (2006). Stress and the suppression of subordinate reproduction in cooperatively breeding meerkats. Proc. Natl. Acad. Sci., 103, 12005–12010.

